# ZCCHC8 is required for the degradation of pervasive transcripts originating from multiple genomic regulatory features

**DOI:** 10.1101/2021.01.29.428898

**Authors:** Joshua W. Collins, Daniel Martin, Genomics and Computational Biology Core, Shaohe Wang, Kenneth M. Yamada

## Abstract

The vast majority of mammalian genomes are transcribed as non-coding RNA in what is referred to as “pervasive transcription.” Recent studies have uncovered various families of non-coding RNA transcribed upstream of transcription start sites. In particular, highly unstable promoter upstream transcripts known as PROMPTs have been shown to be targeted for exosomal degradation by the nuclear exosome targeting complex (NEXT) consisting of the RNA helicase MTR4, the zinc-knuckle scaffold ZCCHC8, and the RNA binding protein RBM7. Here, we report that in addition to its known RNA substrates, ZCCHC8 is required for the targeted degradation of pervasive transcripts produced at CTCF binding sites, open chromatin regions, promoters, promoter flanking regions, and transcription factor binding sites. Additionally, we report that a significant number of RIKEN cDNAs and predicted genes display the hallmarks of PROMPTs and are also substrates for ZCCHC8 and/or NEXT complex regulation suggesting these are unlikely to be functional genes. Our results suggest that ZCCHC8 and/or the NEXT complex may play a larger role in the global regulation of pervasive transcription than previously reported.

## INTRODUCTION

Historically, most transcriptional studies focused on the protein-coding portion of the mouse or human genome under the conventional wisdom that these 20,000 or so genes were interspersed throughout a largely non-transcribed, non-functional genome. Technical advances that brought about genome-wide analyses of transcription unveiled the reality that the vast majority of the genome is, in fact, transcribed as non-protein-coding RNAs, or non-coding RNAs (ncRNAs). Though initially controversial, the scientific community has come to accept the reality of the phenomenon now known as “pervasive transcription” (1-4).

The significance of pervasive transcription is yet to be fully understood and a number of recent studies have described various classes of transcripts of unknown function. Of particular interest to the work presented here are transcripts located upstream of the transcription start site known as promoter upstream transcripts (PROMPTs) (5-7) and upstream antisense RNAs (uaRNAs) (8). Early estimates suggest PROMPTs are ∼200–600 nucleotides in length, highly unstable and short lived, and have both sense and anti-sense orientations (6,7). Similarly short-lived, uaRNAs range from 40– 1,100 bases and are produced predominately at divergently-transcribed, CpG-rich promoters in mouse embryonic stem cells (8). Both PROMPTs and uaRNAs can become 5’ capped, 3’ polyadenylated, and stabilized by exosome depletion (5-8).

The nuclear exosome targeting complex (NEXT) is responsible for exosomal targeting of PROMPTs, enhancer RNAs (eRNAs), 3’ extended small nuclear RNAs (snRNAs), 3’ extended histone RNAs, and intronic RNAs (9-11). NEXT functions in the nucleoplasm and serves as an adapter complex that facilitates recognition and presentation of RNAs to the nuclear exosome (10-12). The trimeric NEXT complex consists of the RNA helicase MTR4/SKIV2L2 which serves to unwind the targeted RNA and directly interacts with the RNA exosome (13,14), a zinc-knuckle scaffold ZCCHC8 and the RNA binding protein RBM7 (10).

Recently, in experiments regarding developing mouse salivary glands and salivary gland cells, we identified ZCCHC8 as having a potentially important function. Here we report a comprehensive and comparative genome-wide analysis of the effects of *Zcchc8* knockout. In addition to its known RNA substrates, we have discovered that ZCCHC8 and/or the NEXT complex are responsible for the targeted degradation of pervasive transcripts produced at CTCF binding sites, open chromatin regions, promoters, promoter flanking regions, and transcription factor binding sites. Further, besides identifying roles in suppressing levels of pervasive, spatially widespread non-coding transcripts, these in-depth analyses reveal that a surprising number of current RIKEN cDNAs and predicted genes appear to be PROMPTs that are unlikely to be functional genes.

## MATERIAL AND METHODS

### Cell Culture and Maintenance

Wild-type, *Btbd7-* and *Zcchc8-*knockout SIMS cells were maintained in phenol red-free DMEM (GE Healthcare/Cytiva, SH30284.01) supplemented with 10% fetal bovine serum (FBS; GE Healthcare/Cytiva, SH30070.03) and incubated at 37°C with 10% CO2. Cells were passaged every three to four days using trypsin-EDTA (Thermo Fisher, 25300120) after rinsing with HBSS (Thermo Fisher, 14170161). Cell density was determined using an automated cell counter (Nexcelom Cellometer Auto 2000).

### CRISPR/Cas9 Knockout of *Zcchc8* in SIMS Cells

The *Zcchc8* KN2.0 non-homology mediated mouse gene knockout kit (KN519669) was purchased from OriGene. Either the pCas-Guide CRISPR vector (OriGene, KN519669G1) containing a single guide RNA target sequence 5’-TAGGTCGCCAAAATCCACAC-3’ or the pCas-Guide CRISPR vector (OriGene, KN519669G2) containing a single guide RNA target sequence 5’-CGAGGCGTTTGACCCACCAG-3’ or a combination of the two was transfected into the mouse submandibular salivary gland cell line SIMS using Thermo Fisher’s Lipofectamine 3000 Reagent kit in the following manner. Approximately 1 × 10^6^SIMS cells were plated in each well of a 6-well plate the day prior to transfection. On the day of transfection, either 3.75 or 7.5 *μ*l of Lipofectamine 3000 reagent was diluted into 125 *μ*l of serum-free Opti-MEM in separate 1.5 ml microfuge tubes for each guide RNA or the combination. In separate 1.5 ml microfuge tubes, 1.0 *μ*g of either pCas-Guide CRISPR vector or the combination, 1.0 *μ*g of linear donor cassette with EF1a promoter followed by eGFP-P2A-Puromycin resistance (OriGene, KN519669D) and 2.0 *μ*l of P3000 reagent per *μ*l DNA was diluted into 125 *μ*l of serum-free Opti-MEM. The diluted DNA mixtures were then added to the respective Lipofectamine microfuge tubes and the reactions were incubated for 15 minutes at room temperature. After incubation, the DNA-lipid mixtures were added drop-wise to the SIMS cells in the respective individual wells. After 48 hours post-transfection, the cells were split 1:10 into DMEM (Thermo Fisher, 11965118) + 10% FBS every three days for a total of 4 passages. The cells were then grown in the selective medium DMEM + 10% FBS + 2 *μ*g/ml puromycin (MilliporeSigma, P8833) for approximately one month. A subset of cells were harvested to check for *Zcchc8* knockout efficiency via western blot, immunofluorescence, and PCR. At this time, it was determined that the greatest knockout efficiency had been achieved using the pCas-Guide CRISPR vector with guide RNA sequence of 5’-TAGGTCGCCAAAATCCACAC-3’ (OriGene, KN519669G1). These cells were chosen to produce individual clones using standard single cell cloning techniques in 96-well plates.

### Western Blots

Approximately 1 × 10^6^ cells from SIMS wild-type, *Btbd7*- and *Zcchc8*-knockout clones were seeded into 10 cm dishes. At ∼75% confluence, cells were washed with pre-chilled PBS followed by the addition of 500 *μ*l of pre-chilled RIPA buffer (25 mM Tris, pH 7.4, 150 mM NaCl, 1.0% NP-40, 0.5% sodium deoxycholate, 0.1% SDS) supplemented with 1× Halt Protease and Phosphatase Inhibitor Cocktail (Thermo Fisher, 78444). Cells were scraped on ice and collected in pre-chilled 1.5 ml tubes (Eppendorf, 022363212). The cell suspensions were incubated on ice for 30 minutes followed by centrifugation at 13,000 rpm for 15 min at 4°C. Supernatants were transferred to pre-chilled 1.5 ml tubes and stored at -20°C. Lysates were quantified using the Pierce BCA Protein Assay Kit (Thermo Fisher, 23227). Aliquots of 25 *μ*g lysate were denatured in 1X Laemmli sample buffer (Bio-Rad, 1610747) at 99°C for 5 min. Using a Bio-Rad Mini-PROTEAN Tetra Vertical Electrophoresis Cell, lysates and 5 *μ*l of Precision Plus Protein Kaleidoscope Standards (Bio-Rad, 1610375) were run on Bio-Rad 7.5% Mini-PROTEAN TGX precast gels with Tris/Glycine/SDS (Bio-Rad, 1610732) running buffer at 115 V followed by transfer to Bio-Rad Trans-Blot Turbo 0.2 %m nitrocellulose via a Bio-Rad Trans-Blot Turbo Transfer system. Membranes were then incubated in blocking solution consisting of 5% nonfat dry milk in TBST (Tris Buffered Saline with 0.5% Tween-20; Quality Biological, 351-086-101; MilliporeSigma, P2287) for 1 hour at room temperature followed by incubation with primary antibodies diluted in blocking solution overnight at 4°C. Membranes were washed 3 times for 15 min each in TBST and incubated with LI-COR secondary antibodies diluted in Blocking Solution for 1 hour at room temperature protected from light. The membranes were then washed 3 times for 15 min each in TBST at room temperature and imaged on a LI-COR Odyssey CLx imaging system controlled by the LI-COR Image Studio software. The primary antibody dilutions were 1:10,000 anti-Zcchc8 (Proteintech, 23374-1-AP), 1:10,000 anti-*α*-tubulin (MilliporeSigma, T6199). The secondary antibody dilutions were 1:10,000 680RD goat anti-mouse (LI-COR, 926-68070) and 1:10,000 800CW goat anti-rabbit (LI-COR, 926-32211).

### Immunofluorescence and Confocal Microscopy

Approximately 1 × 10^5^cells from SIMS wild-type, *Btbd7*- and *Zcchc8*-knockout clones were seeded into 35 mm MatTek dishes (MatTek, P35G-1.5-20-C) and grown for 48 hours. Cell medium was removed, and the cells were fixed with 2 ml of 4% PFA in PBS (Electron Microscopy Sciences, 15710) for 20 min at room temperature. Cells were then quickly washed with 2 ml of PBS followed by permeabilization with 1 ml of PBS with 0.1% Triton X-100 (Thermo Fisher, 28314) for 10 min at room temperature. Cells were then washed with 2 ml of wash buffer (PBS + 0.5% Tween-20) and then blocked for 1 hour with 1 ml of blocking buffer (wash buffer containing 3% fatty acid free BSA (Thermo Fisher, 126609)). The cells were then incubated with 200 *μ*l of primary antibodies diluted in blocking buffer overnight at 4°C. The cells were then washed with 1 ml of wash buffer three times for 5 min each and incubated with 100 *μ*l of secondary antibodies diluted in blocking buffer for 1 hour at room temperature. Cells were then washed with 1 ml of wash buffer three times for 5 min each. After the final wash, 12 *μ*l of Fluoro-Gel II with DAPI mounting medium (Electron Microscopy Sciences, 17985-50) was added to the cells on the glass surface of the MatTek dish. A coverslip was then sealed over the glass surface to protect the cells from damage. The cells were then imaged using either a 40× C-Apochromat, 1.1 NA or 63× C-Apochromat, 1.2 NA objective on a Zeiss LSM 880 system controlled by Zeiss ZEN software. The primary antibody dilution was 1:500 anti-Zcchc8 (Proteintech, 23374-1-AP). The secondary antibody dilution was 1:200 Alexa Fluor 647 donkey anti-rabbit (Jackson ImmunoResearch, 711-606-152). Additionally, cells were stained with a 1:200 dilution of rhodamine Phalloidin (Thermo Fisher, R415) during the secondary antibody incubation step.

### RNA Extraction and Sequencing Library Preparation

Approximately 1 × 10^5^ cells from SIMS wild-type, *Btbd7*- and *Zcchc8*-knockout clones were seeded into 60 mm dishes and grown until cells reached ∼75% confluence. Cell medium was removed and 0.5 ml of TRIzol Reagent (Thermo Fisher, 15596026) was added directly to the dishes. After brief trituration, the lysates were collected into 1.5 ml tubes and incubated for 5 min at room temperature. Next, 0.1 ml of chloroform was added to each lysate and mixed thoroughly by inverted the tubes multiple times. The lysates were incubated at room temperature for three minutes and then centrifuged at 13,000 rpm for 15 min at 4°C. The aqueous phase of each lysate was then transferred to separate, fresh 1.5 ml tubes and 0.25 ml of 70% ethanol was added to each tube. The samples were then transferred to RNeasy spin columns and 2 ml collection tubes from an RNeasy Mini Kit (Qiagen, 74104) and centrifuged at 13,000 rpm for 15 sec. The flow-through was discarded and on-column DNase digestion was performed using the RNase-Free DNase Set (Qiagen, 79254) and the following protocol from Qiagen: 350 *μ*l of buffer RW1 was added to each column followed by centrifugation at 13,000 rpm for 15 sec. The flow-through was discarded. 80 *μ*l of DNase I incubation mix (10 *μ*l DNase I stock solution in 70 *μ*l buffer RDD) was added to each column membrane and incubated for 15 min at room temperature. 350 *μ*l of buffer RW1 was added to each column followed by centrifugation at 13,000 rpm for 15 sec. The flow-through was discarded and 500 *μ*l of buffer RPE was added to each column followed by centrifugation at 13,000 rpm for 15 sec. Again, 500 *μ*l of buffer RPE was added to each column followed by centrifugation at 13,000 rpm for 2 min. The RNeasy columns were then placed in new 2 ml collection tubes and centrifuged for 13,000 rpm for 1 min. The RNeasy spin columns were then placed in new 1.5 ml collection tubes and 50 *μ*l of RNase-free water was added to the column membranes followed by centrifugation at 13,000 rpm for 1 min. The RNA samples were then stored at -80°C.

RNA integrity was assessed using a Fragment Analyzer (Agilent) and sequencing libraries were prepared using the Illumina TruSeq Stranded Total RNA library preparation kit according to the manufacturer’s protocols. Library preparation was performed on four replicates of each cell line. The samples were analyzed on an Illumina NextSeq500 configured for 40 paired-end reads and the quality of the reads was evaluated using FastQC software.

### Read Mapping and Differential Expression Analysis

In order to perform read mapping of PROMPTs and genomic regulatory features, the respective gtf/gff files were revised in order to generate the genome files for STAR read alignment. The Ensembl regulatory build gff file was filtered for each genomic regulatory feature and a new gff file was created for each. Each genomic feature was given a specific identifier in the feature identifier column of the respective new regulatory build gff files (ctcf, enhancer, ocr, promoter, pfr, tfbs). STAR genome generation was then conducted with the additional parameters --sjdbGTFfeatureExon feature and --sjdbGTFtagExonParentGene ID.

For PROMPTs, the primary assembly gtf file was filtered to remove all feature types except for the “gene” identifier. This identifier was changed to “prompt” and the genomic coordinates were altered to 3 kb upstream of the start site. STAR genome generation was then conducted with the additional parameters --sjdbGTFfeatureExon prompt and --sjdbGTFtagExonParentGene ID.

Read mapping against the recent GENCODE mouse release M24 was performed using the STAR 2.7.3a aligner using the standard mode with mapping parameters derived from the GENCODE project. Additionally, read counting was performed using the quantMode utility of STAR.

Read counts were filtered to remove low expressing features (genes, PROMPTs, genomic regulatory features, etc., with <5 counts in at least one sample). Differential expression was evaluated by three independent statistical methods (DESeq2, edgeR and Limma-Voom).

### Bioinformatics and Data Analysis

Data analysis was performed using the SAMtools (15), deepTools (16), PhenoGram (17), and R software packages. Scripts used for data analysis can be found at https://github.com/collinsjw/Zcchc8-KO-pervasive-transcripts.

### Data Availability

RNA sequencing data is deposited in NCBI GEO (accession number GSE165689).

## RESULTS

### *Zcchc8* knockout in mouse salivary gland cells

In order to test the hypothesis that ZCCHC8 has some additional function besides its known scaffolding role in the NEXT complex, we used the CRISPR/Cas9 system to knockout *Zcchc8* in the SIMS mouse salivary gland cell line (Figure 1A). We then isolated >20 clonal populations using standard single cell cloning techniques. After western blotting for ZCCHC8, we randomly selected three individual cell lines from those with depleted ZCCHC8 expression. Genetic sequencing confirmed differing two-base-pair deletions near the CRISPR/Cas9 target sequence that resulted in newly formed stop codons shortly downstream from the deletion site in each clone (Figure 1A). Follow-up western blotting and immunofluorescence analyses confirmed the absence of ZCCHC8 protein expression in the selected clones (Figure 1B-C). Gross morphological examination did not reveal any noticeable differences between the control and deletion cell lines.

**Figure 1.**
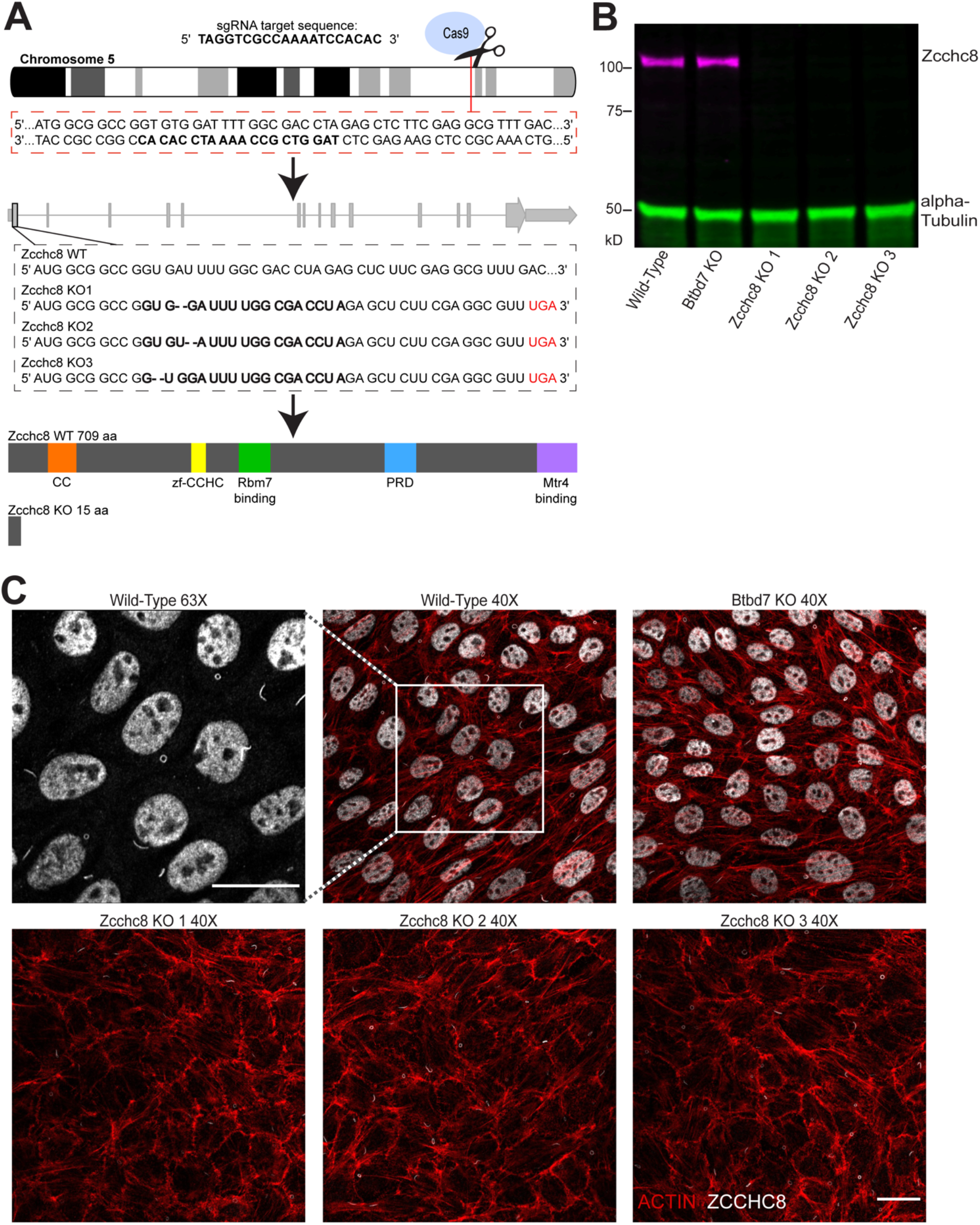
(A) Schematic depicting the CRISPR/Cas9 strategy for disrupting *Zcchc8* in SIMS mouse salivary gland cells. Single cell cloning produced three separate knockout clones with two-base-pair deletions ∼12-15 bp downstream of the TSS. The resulting frameshift produced a translational stop codon at amino acid position 16. (B) Western blot confirmation of ZCCHC8 ablation in three separate clones. Western blots were probed with a rabbit, polyclonal anti-ZCCHC8 antibody from Proteintech and a mouse, monoclonal anti-α-Tubulin antibody from Sigma. The composite image of the blot was produced using a LI-COR Odyssey CLx imaging system controlled by the LI-COR Image Studio software. (C) Immunofluorescence confirmation of ZCCHC8 ablation in three separate clones compared to wild-type and knockout of the unrelated gene *Btbd7* (negative control). The ZCCHC8 immunofluorescence (greyscale) dynamic range was equal for all images. The same anti-ZCCHC8 antibody as in (B) was used. Note the presence and loss of nuclear ZCCHC8 staining compared to the continued presence of antibody cross-reactivity to primary cilia before and after *Zcchc8* knockout. Actin fibers (red) were stained with a rhodamine-labeled phalloidin from Thermo Fisher. Actin images were processed using a rolling-ball background subtraction method. Scale bar = 20 μm.

We then performed a series of biological assays to assess the effects of *Zcchc8* knockout in this cell line. Somewhat remarkably, these *Zcchc8* knockout clones showed negligible differences in MTT proliferation, Matrigel invasion, colony formation, spheroid formation, soft agar, and scratch assays when compared to wild-type control (data not shown).

### RNA sequencing of *Zcchc8* knockout cells

In the absence of any observed in vitro biological phenotype, we proceeded to perform RNA sequencing on wild-type and *Zcchc8* knockout SIMS cells. As an additional negative control, we also included SIMS cells in which we previously disrupted the gene *Btbd7* using the CRISPR/Cas9 system. Considering the role of ZCCHC8 and the NEXT complex in RNA degradation pathways, we used the Illumina TruSeq Stranded Total RNA library preparation kit in order to ultimately detect differential expression of small RNAs, lncRNAs, and mRNAs. Library preparation was performed on four replicates of each cell line.

We then performed differential expression analysis of genomic features using three independent statistical tests to assess differential expression between groups: DESeq2, edgeR, and Limma-Voom. These methods differ in normalization of feature expression across samples, assumptions about the distribution of the underlying data and statistical test used, and are widely accepted in the current literature.

Principal component analysis (PCA), multidimensional scaling (MDS), and Euclidean distance clustering (EDC) of gene expression data revealed a single wild-type SIMS replicate that differed significantly from all of the other wild-type replicates (Supplemental Figures S1A and S2A). This replicate was removed from all subsequent analyses. The replicates of all other clonal groups were highly similar and were included in all analyses (Figure 2A-B, Supplemental Figures S1B and S2B). Importantly, unsupervised EDC revealed two major clusters: a control cluster consisting of wild-type and Btbd7 knockout cells and a *Zcchc8* knockout cluster (Figure 2B). Further, the replicates of each clone clustered together into related subclusters (Figure 2B).

**Figure 2.**
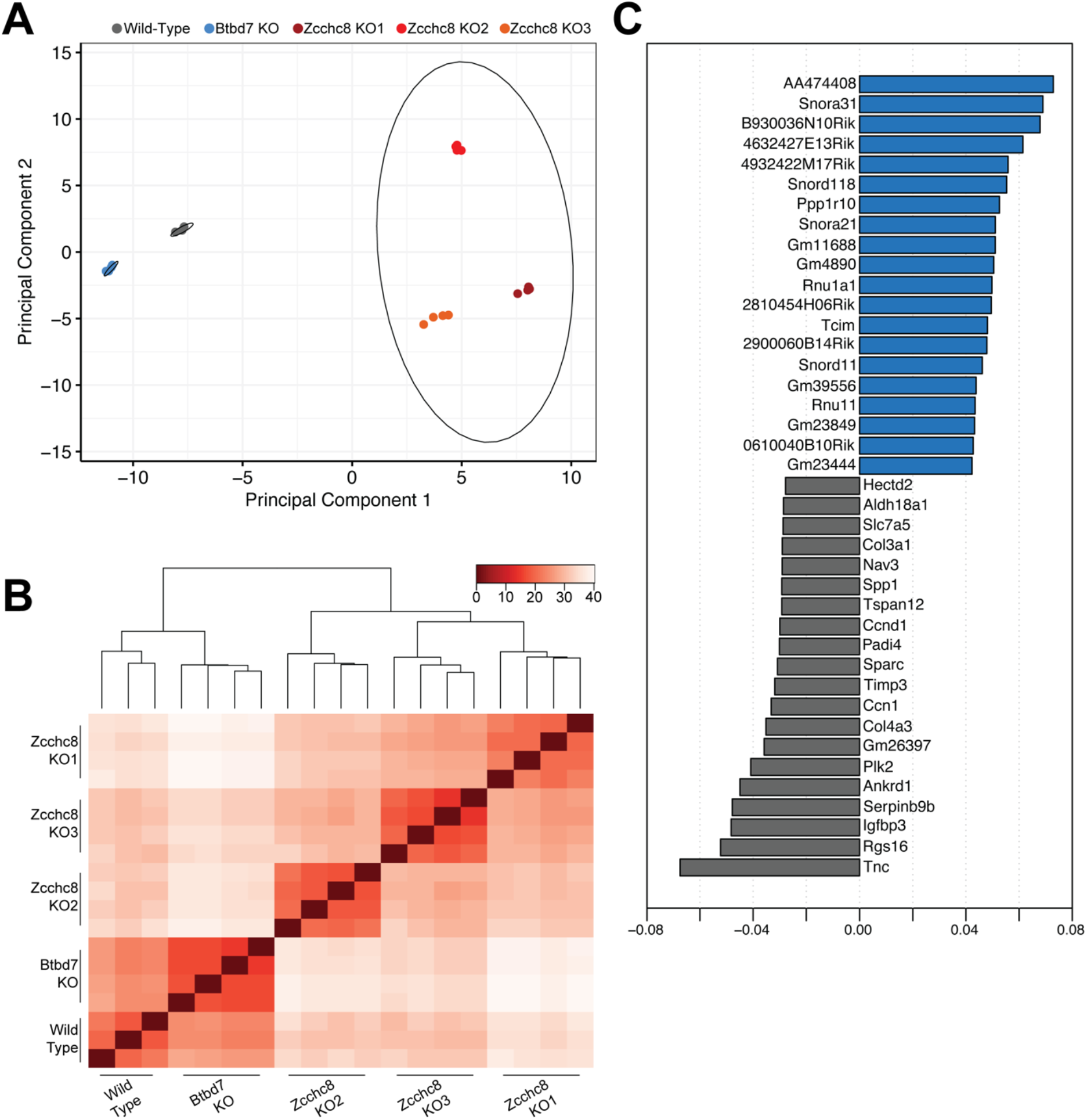
(A) PCA plot of components 1 and 2 using computed variance-regularized and log_2_-transformed gene expression data from DESeq2 analysis. Principal component 1 = 57.35% variance; Principal component 2 = 20.76% variance. Ellipses represent 95% confidence regions. (B) Unsupervised Euclidean Distance Clustering of transformed data from (A). (C) Principal component loadings plot depicting the top 20 genes positively and negatively correlated with principal component 1.

Strikingly, a plot of the top 20 genes both positively and negatively correlated with the variance of the first principal component revealed a large number of RIKEN cDNAs and predicted genes (with the Mouse Genome Informatics naming convention of Gm*xxxxx*) positively correlated with this variance (Figure 2C). In fact, of the top 100 genes positively correlated with the variance of the first principal component, 62 are RIKEN cDNAs and predicted genes (29 and 33, respectively) (Table 1). Only 11 are snRNAs or snoRNAs—two RNA species that are known substrates for ZCCHC8 and the NEXT complex.

**Table 1.**
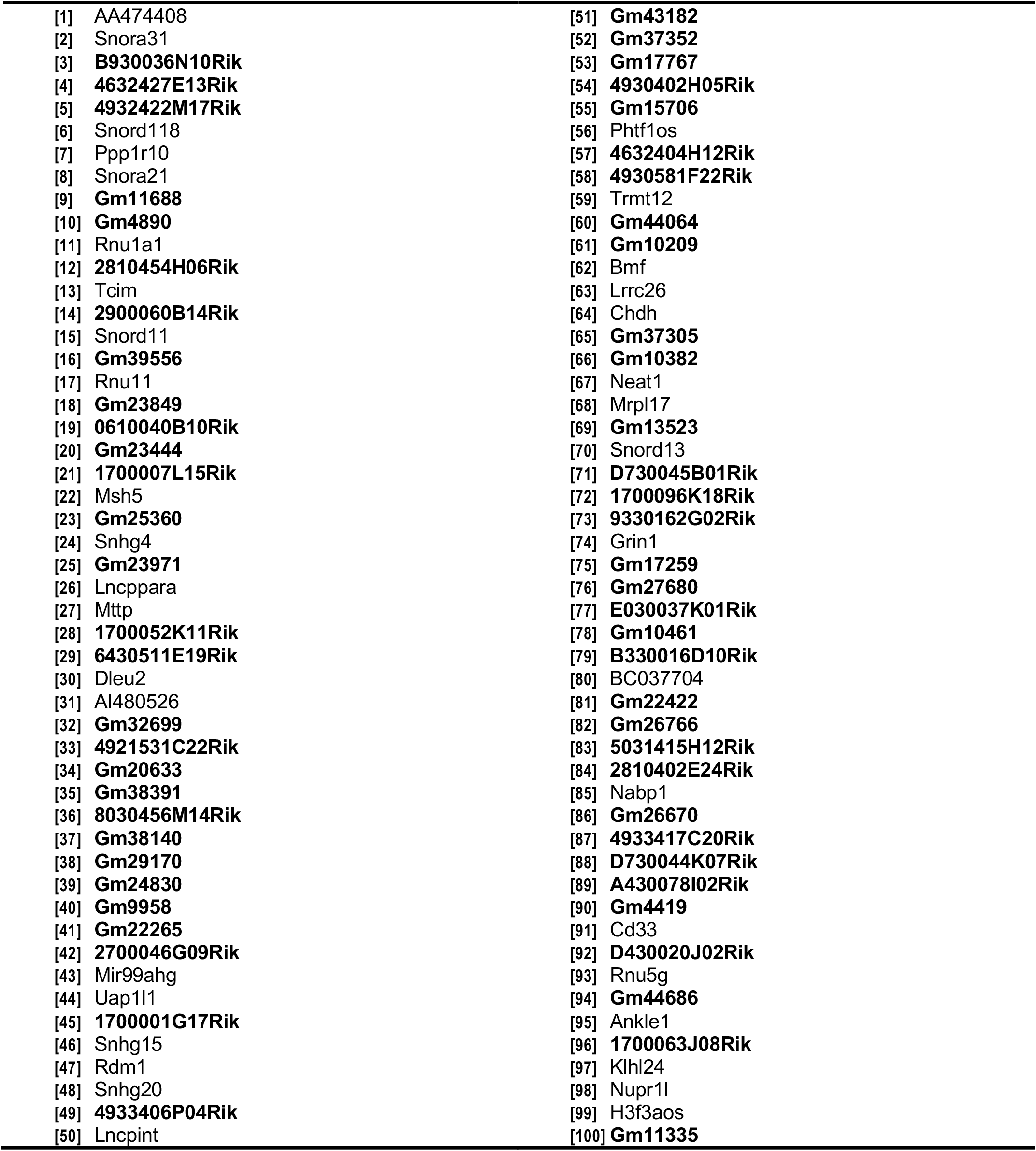
Top 100 genes positively correlated with the variance of the first principal component in order of variance contribution. RIKEN cDNAs and predicted genes are in bold.

### Effects of *Zcchc8* knockout on pervasive transcripts associated with genomic regulatory features

In order to assess the role of ZCCHC8 in the degradation of PROMPT RNAs, we compared the read coverage of the 3 kb region upstream of the TSS of all expressed genes in wild-type and *Zcchc8* knockout cells. As expected, knockout of *Zcchc8* resulted in the appearance of PROMPTs upstream of the TSS for numerous genes (Figure 3A-B). Metagene plots revealed the majority of PROMPTs to be less than 1.5 kb upstream of the TSS (Figure 3A). Read coverage heat maps for all PROMPT regions are shown in Figure 3B. The negative control Btbd7 knockout had no effect on transcription upstream of TSSs (Figure 3A-C).

**Figure 3.**
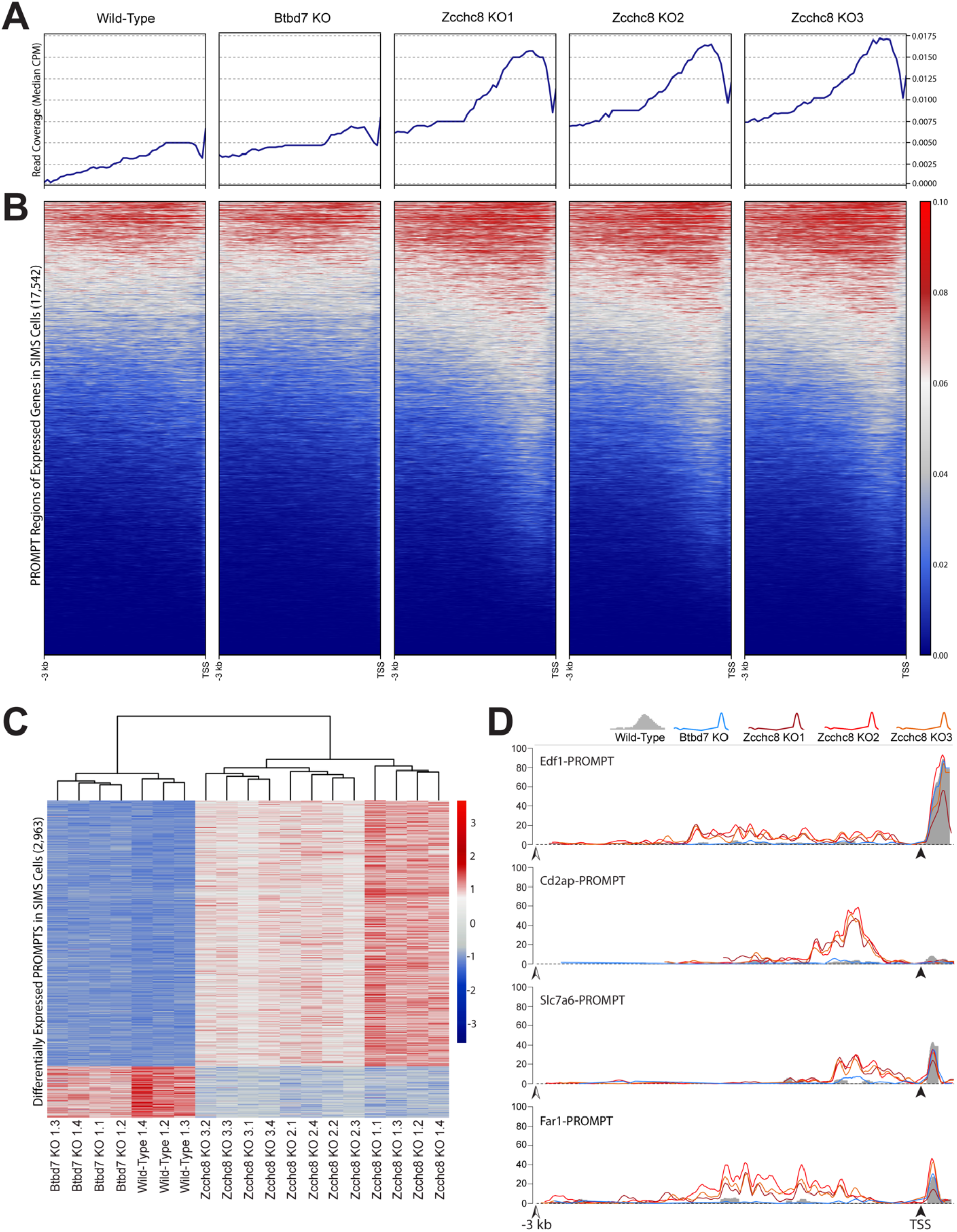
(A) Metagene plots showing the median read coverage for CPM-normalized read counts in 3-kb regions upstream of the TSS for all expressed genes. Genes were considered to be expressed if there were >5 reads in at least one sample. (B) Read coverage heatmaps of the 3-kb upstream regions in (A). In order to create a single plot of all clonal replicates in (A) and (B), read counts from replicate bam files were merged and indexed using the SAMtools software package followed by CPM-normalization. (C) Heatmap using k-means clustering (k = 2) of the 2,963 differentially expressed PROMPTs in SIMS *Zcchc8* knockout cells that are found in common to each of the statistical analysis methods DESeq2, edgeR, and Limma-Voom. (D) Read coverage plots indicating the differential expression of PROMPTs within the 3 kb upstream region of the TSS for the genes *Edf1, Cd2ap, Slc7a6*, and *Far1*. In order to create a single plot of all clonal replicates, read counts from replicate bam files were merged and indexed using the SAMtools software package followed by CPM-normalization prior to plotting using the Gviz R package. Black arrowheads indicate the TSS for each gene followed by the first 250 bp of transcription coverage. Open arrowheads indicate the -3 kb position relative to the TSS.

We next performed differential expression analysis of all PROMPT regions 3 kb upstream of annotated gene start sites. Using cut-off values of >1.5-fold expression difference and <0.01 adjusted p-value, DESeq2, edgeR, and Limma-Voom revealed the presence of 5,393, 5,709, and 4,963 differentially expressed PROMPT regions, with 2,963 in common (Figure 3C, Supplemental Figure S3A, Supplemental Table S1). Hereafter, we will only refer to those differentially expressed PROMPTs, genes, regulatory features, etc., that are shared according to all three statistical methods. Read coverage plots for selected differentially expressed PROMPTs were generated using the Gviz package in R and are shown in Figure 3D. These detailed plots reveal the remarkable similarity of PROMPT transcription read coverage in three separate knockout clones as compared to wild-type and the *Btbd7* knockout negative control (Figure 3D).

Considering that divergent transcription at TSSs is presumed to occur at most active protein coding genes (5,7,18-21), we expected a large number of PROMPTs in our *Zcchc8* knockout cells. Recently, it was shown that the poly(A) exosome-targeting (PAXT) connection can act as a failsafe in the case of NEXT disruption (11). We wondered if NEXT complex redundancy through the PAXT connection would be evident through increased transcription or if the remaining NEXT complex components or RNA exosome were up-regulated as a compensatory mechanism for loss of ZCCHC8 function. Upon further examination, we found no evidence of such regulation at the transcriptional level (Supplemental Table S2). We also mapped the chromosomal distribution of PROMPTs and found they were widespread and well-distributed across all chromosomes (Supplemental Figure S4).

Differential expression analysis also detected ∼15% of PROMPT regions that were down-regulated in *Zcchc8* knockout cells (Figure 3C). As the function of the NEXT complex is to degrade PROMPTs, we examined how ZCCHC8 disruption could result in decreased transcription in these regions. Deeper examination revealed portions of full-length expressed genes overlapping these PROMPT regions in either head-to-head or head-to-tail orientation. Thus, these regional transcriptional differences are due to differential gene expression rather than to authentic PROMPT expression. Hereafter, we will refer to PROMPTs as those 3 kb upstream regions that are up-regulated in *Zcchc8* knockout cells. Accordingly, we found 2,486 PROMPTs in our SIMS *Zcchc8* knockout cells.

Similar to our PROMPT analysis, we performed differential expression analyses on regulatory features (as annotated in the Ensembl Regulation database) and found ZCCHC8 ablation resulted in transcriptional up-regulation of 247 CTCF binding sites, 243 enhancers, 142 open chromatin regions, 104 promoters, 471 promoter flanking regions, and 45 transcription factor binding sites (>1.5-fold expression difference and <0.01 adjusted p-value) (Figure 4A). In negative control Btbd7 knockout cells, there were fewer than 5 differentially expressed regulatory features of each category. The results of this differential expression analysis are summarized in Supplemental Table S1. We mapped the chromosomal locations of these regulatory regions and found them to be well-distributed along each chromosome with the exception of chromosome X (Figure 4B). Further, we evaluated whether any of these regulatory features overlapped with PROMPTs and found that only a small number shared intersecting genomic coordinates (Figure 4C).

**Figure 4.**
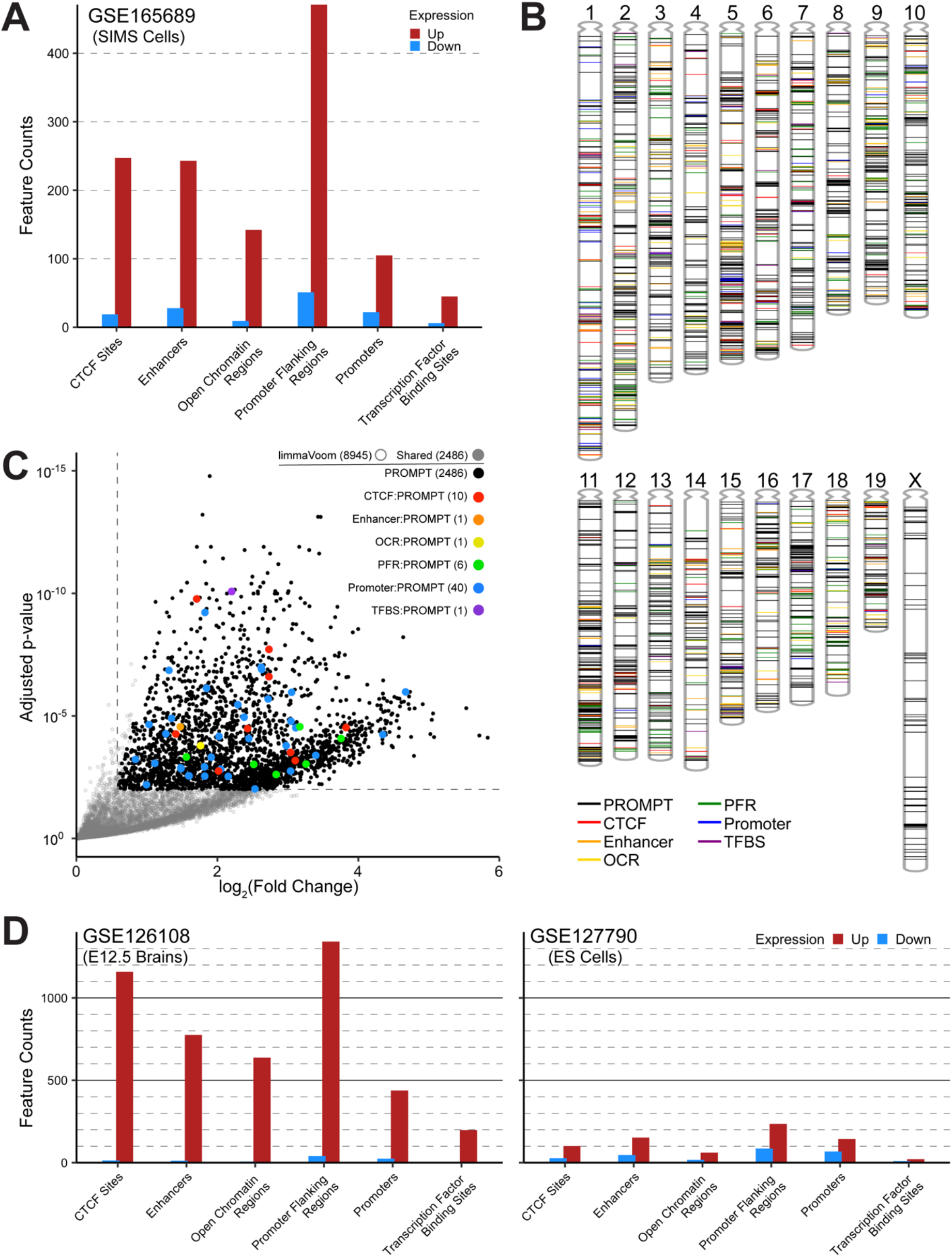
(A) Bar chart showing the number of differentially expressed genomic regulatory features in SIMS *Zcchc8* knockout cells. (B) Chromosomal distribution of differentially expressed PROMPTs and genomic regulatory features from SIMS *Zcchc8* knockout cells (GSE165689). (C) Scatter plot of PROMPTs and genomic regulatory features with overlapping genomic coordinates. Data were generated using the LimmaVoom statistical analysis. Horizontal and vertical dashed lines demarcate adjusted p-value of 0.01 and fold-change of 1.5 (log_2_(1.5) ≈ 0.584), respectively. Open circles indicate those PROMPTs that are specific to the LimmaVoom analysis. Closed circles indicate PROMPTs that are shared within DESeq2, edgeR, and LimmaVoom analyses and meet the significance thresholds of >1.5 fold-change and <0.01 adjusted p-value. Colored circles indicate genomic regulatory features with overlapping genomic coordinates to PROMPTs. (D) Bar chart showing the number of differentially expressed genomic regulatory features found in E12.5 brains (GSE126108) and ES cells (GSE127790) from *Zcchc8* knockout mice.

### Differential gene expression in *Zcchc8* knockout cells

We next turned our attention to differential gene expression using the same analyses we used for PROMPTs. Using the same thresholds of >1.5-fold expression change and adjusted p-value of <0.01, we found a total of 852 differentially expressed genes in our SIMS cells (Figure 5A, Supplemental Figure S3B). Of these genes, 712 were up-regulated in *Zcchc8* knockout cells while only 140 were down-regulated (Figure 5A, Supplemental Table S1). Intriguingly, of the 712 up-regulated genes in the *Zcchc8* knockout cells, 486 were RIKEN cDNAs or predicted genes (Figure 5B) affirming our exploratory PCA findings. Only 9 down-regulated RIKEN cDNAs and predicted genes were discovered (Figure 5B). Conversely, the Btbd7 knockout cells (negative control) showed only 62 up-regulated and 80 down-regulated genes with only 2 and 8 of these consisting of RIKEN cDNAs or predicted genes, respectively (Supplemental Figure S5).

**Figure 5.**
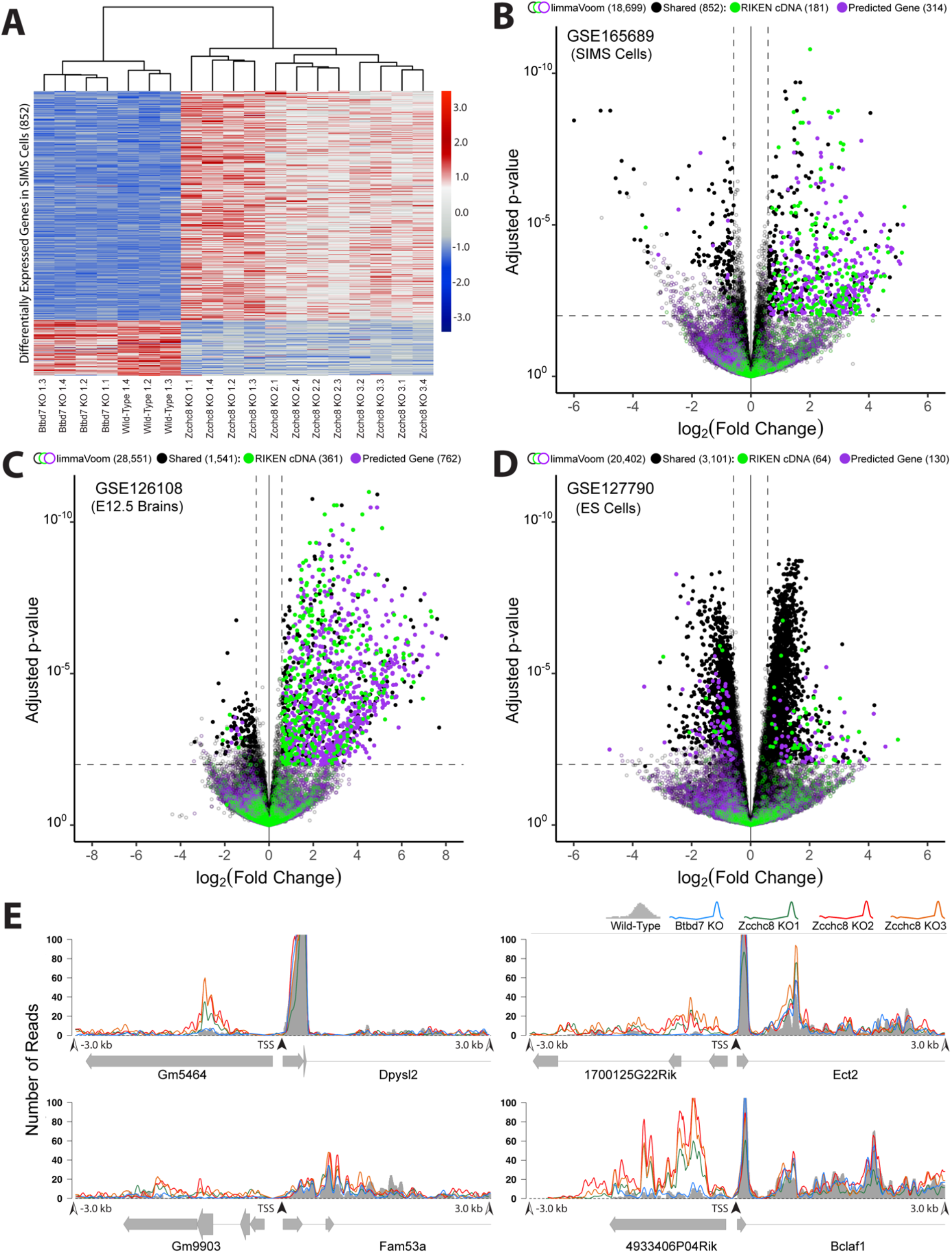
(A) Heatmap using k-means clustering (k = 2) of the 852 differentially expressed genes in SIMS *Zcchc8* knockout cells that are found in common to each of the statistical analysis methods DESeq2, edgeR, and Limma-Voom. (B) Scatter plot of gene expression data in (GSE165689) SIMS *Zcchc8* knockout cells. Data were generated using the LimmaVoom statistical analysis. Horizontal and vertical dashed lines demarcate adjusted p-value of 0.01 and fold-change of 1.5 (log_2_(1.5) ≈ 0.584), respectively. Open circles indicate those genes that are specific to the LimmaVoom analysis. Closed circles indicate genes that are shared within DESeq2, edgeR, and LimmaVoom analyses and meet the significance thresholds of >1.5 fold-change and <0.01 adjusted p-value. Colored circles indicate RIKEN cDNAs (green) and predicted genes (magenta). (C) Scatter plot of gene expression data (GSE126108) in E12.5 brains taken from *Zcchc8* knockout mice as in (A). (D) Scatter plot of gene expression data (GSE127790) in ES cells taken from *Zcchc8* knockout mice as in (A). (E) Read coverage plots indicating the differential expression of RIKEN cDNAs and predicted genes upstream of protein coding genes. In order to create a single plot of all clonal replicates, read counts from replicate bam files were merged and indexed using the SAMtools software package followed by CPM-normalization prior to plotting using the Gviz R package. Black arrowheads indicate the TSS for each protein coding gene. Open arrowheads indicate the +/-3 kb position relative to the TSS. Note that the full length of the RIKEN cDNAs and predicted genes are shown while only the first 3 kb of the protein coding genes are shown. Read coverage y-axes were truncated at 100 reads in order to maintain scale.

This unexpected finding stimulated us to examine the relationship between these RIKEN cDNAs and predicted genes and the genomic regulatory features governed by ZCCHC8 and the NEXT complex. Surprisingly, 334 out of the 486 up-regulated RIKEN cDNAs and predicted genes had intersecting genomic coordinates with the differentially expressed PROMPTs in our data set. Read coverage plots for selected RIKEN cDNAs and predicted genes revealed expression patterns that display the trademarks of PROMPTs (Figure 5E).

Lastly, we asked if ZCCHC8 and/or the NEXT complex might directly regulate specific genes or pathways. We subjected the remaining differentially expressed genes to Signaling Pathway Impact Analysis (SPIA), PathNet, and Ingenuity Pathway Analysis in order to generate candidate genes and pathways. Our efforts at comparing these candidates did not reveal any selective effects on the pathways we singled out for validation.

### Effects of *Zcchc8* knockout on pervasive transcripts associated with genomic regulatory features in mice

Two other research groups recently developed *Zcchc8* knockout mice (22,23). Gable et al., performed RNA-seq on E12.5 brains while Wu et al., performed RNA-seq on ES cells derived from knockout mice. Using the publicly available raw data (NCBI GEO: GSE126108 and GSE127790) from these experiments, we performed our same differential expression analyses to assess the global effects of *Zcchc8* disruption.

Interestingly, we found 4,740 PROMPTs in E12.5 brains and only 422 PROMPTs in ES cells. We found our SIMS cells had 1,772 PROMPTs in common with E12.5 brains and only 138 PROMPTs in common with ES cells. There were 132 PROMPTs common to all three groups (Supplemental Table S3).

Considering our establishment of the role of ZCCHC8 and/or the NEXT complex in regulating pervasive transcription of genomic regulatory features in SIMS *Zcchc8* knockout cells, we asked if our findings would be recapitulated in the knockout mouse datasets. Indeed, in E12.5 brains (GSE12618) we found ZCCHC8 ablation resulted in the transcriptional up-regulation of 1,159 CTCF binding sites, 775 enhancers, 638 open chromatin regions, 438 promoters, 1,342 promoter flanking regions, and 198 transcription factor binding sites (>1.5-fold expression difference and <0.01 adjusted p-value) (Figure 4D). Likewise, in ES cells (GSE127790) we found 101 CTCF binding sites, 152 enhancers, 61 open chromatin regions, 143 promoters, 235 promoter flanking regions, and 21 transcription factor binding sites (>1.5-fold expression difference and <0.01 adjusted p-value) (Figure 4D). The number of genomic regulatory features shared between all three data sets are summarized in Supplemental Table S3.

We further explored if our discovery of the relationship between PROMPTs, RIKEN cDNAs and predicted genes would also be recapitulated in the knockout mouse datasets. First, we found 1,541 differentially expressed genes in E12.5 brains with 1,410 up-regulated and only 131 down-regulated in knockouts (Figure 5C). We found 3,101 differentially expressed genes in ES cells with 1,804 up-regulated and 1,297 down-regulated (Figure 5D). In comparison, our SIMS cells had 450 up-regulated genes and only 1 down-regulated gene in common with E12.5 brains; 89 up-regulated genes and only 7 down-regulated genes in common with ES cells. Among the three knockout groups, there were 69 common up-regulated genes and 0 down-regulated genes in common. These data are summarized in Supplemental Table S3.

Once again, closer inspection revealed a large proportion of the up-regulated genes in the *Zcchc8* knockout groups were RIKEN cDNAs and predicted genes. Remarkably, of the 1,410 up-regulated genes in E12.5 brains, 1,123 are RIKEN cDNAs or predicted genes (Figure 5C) with 645 having overlapping genomic coordinates with PROMPTs. Further, of the 450 common up-regulated genes between our SIMS cells and the E12.5 brains, 372 were RIKEN cDNAs or predicted genes. In an intriguing contrast, only 102 of the 1,804 up-regulated genes in ES cells were RIKEN cDNAs or predicted genes (Figure 5D). These data are summarized in Supplemental Table S3.

## DISCUSSION

This study provides a comprehensive examination of the effects of ZCCHC8 ablation on the mouse transcriptome. Our results show that ZCCHC8 and/or the NEXT complex regulates a larger family of ncRNA species than previously reported. We show that disruption of *Zcchc8* results in the up-regulation of transcriptional products in multiple genomic regulatory regions including CTCF binding sites, enhancers, open chromatin regions, promoters, promoter flanking regions, and transcription factor binding sites. Further, we have shown >95% of these genomic regions have non-overlapping genomic coordinates with PROMPTs in our SIMS *Zcchc8* knockout cells. Thus, we conclude that, in addition to established substrates, ZCCHC8 and/or the NEXT complex serve to regulate pervasive transcription at CTCF binding sites, open chromatin regions, promoters, promoter flanking regions, and transcription factor binding sites. To our knowledge, this is the first such comprehensive report.

Surprisingly, we found ZCCHC8 ablation resulted in a large increase in the expression of RIKEN cDNAs and predicted genes. Given the transcriptional coverage, profile, and orientation of these transcripts, our results suggest that a significant number of RIKEN cDNAs and predicted genes are, in fact, PROMPTs (though ultimately experimental confirmation will be required). Using rather stringent criteria, we found 334 RIKEN cDNAs and predicted genes in our SIMS cells, and 645 in E12.5 brains, that display the hallmarks of PROMPTs. Considering that ZCCHC8 and the NEXT complex primarily serve to regulate pervasive transcription of a large number of ncRNA families, it stands to reason that many of the remaining up-regulated RIKEN cDNAs and predicted genes that are non-overlapping with PROMPTs may also be pervasive transcripts from other ncRNA families. Less likely, the possibility remains that many of these RIKEN cDNAs and predicted genes constitute a specific set of protein-coding genes that are negatively regulated in large part by ZCCHC8 and/or the NEXT complex.

As expected, the overall genomic expression profile significantly differed between SIMS cells, E12.5 brains, and ES cells. Interestingly, despite the obvious difference between mouse salivary glands cells and E12.5 brains, these groups shared 1,772 PROMPTs and 372 predicted genes and RIKEN cDNAs while relatively little overlap exists with these groups and ES cells. Additionally, our SIMS cells and the E12.5 brains had ∼5.9 and 11.2 times more PROMPTs, respectively, and ∼4.8 and 11.0 times more up-regulated RIKEN cDNAs and predicted genes, respectively, than ES cells.

Obviously, these results are indicative of the larger transcriptomic differences among cells of varying differentiation potential and suggest that epigenomic and heterochromatic organization of the genome during these states may play a role in PROMPT expression and/or degradation. Considering that it is generally accepted that ES cells have more open, plastic chromatin with reduced nucleosome density than differentiated cells, it is seemingly counterintuitive that the more differentiated state of SIMS cells and E12.5 brains may better lend itself to conditions suitable for increased transcription of PROMPTs. Nevertheless, recent research using a highly sensitive chemical mapping of nucleosome organization in mouse ES cells has shown that, contrary to the prevailing model, nearly all genes have a class of “fragile” nucleosomes occupying previously designated nucleosome-depleted regions upstream of transcription start sites (24). Further, this nucleosome mapping showed a high degree of nucleosome occupancy at CTCF sites (24). Though this finding is currently considered controversial, it has clear implications for potentially suppressing PROMPT and genomic regulatory feature transcription in ES cells.

Alternatively, it is speculated that as RNA polymerases elongate the sense transcript, negative supercoiling of the DNA upstream of the TSS can prime antisense transcription initiation (20). Perhaps, the epigenomic and heterochromatic landscape of more differentiated cells is more conducive to negative supercoiling that would prime such antisense transcription in a narrow window upstream of the TSS.

Regardless of the mechanism, it is clear that ES cells have fewer PROMPTs than our SIMS cells and the E12.5 brains. Perhaps this should not come as a surprise considering the potential detrimental effects that pervasive transcripts, like PROMPTs, may have on the tight regulation required to maintain stemness. It would also not be surprising if future research were to uncover a more generalized, global regulatory system for preventing these unwanted transcripts in ES cells. To potentially shed light on the mechanism, it would be interesting to profile the DNA methylation state, nucleosome density, and nucleosome modification status of the PROMPT regions shared in SIMS cells and E12.5 brains versus the ES cells.

## AVAILABILITY

Scripts used for data analysis can be found at: https://github.com/collinsjw/Zcchc8-KO-pervasive-transcripts.

## ACCESSION NUMBERS

RNA sequencing data is deposited in NCBI GEO (accession number GSE165689).

## Supporting information

Graphical Abstract

## ACKNOWLEDGEMENT

We thank all the members of the Yamada lab for helpful discussions and critical reading of the manuscript and A.D. Doyle from the NIDCR Imaging Core for microscopy assistance. This work utilized the computational resources of the NIH HPC Biowulf cluster.

Genomics and Computational Biology Core

Robert J. Morell^1^, Erich T. Boger^1^, Daniel Martin^2^, Kerianne Richards^1^, Zheng Wei^2^

^1^National Institute on Deafness and Other Communication Disorders, National Institutes of Health, Bethesda, Maryland, USA.

^2^National Institute of Dental and Craniofacial Research, National Institutes of Health, Bethesda, Maryland, USA.

## AUTHOR CONTRIBUTIONS

J.W.C. and K.M.Y. conceived the project. J.W.C. designed and conducted the experiments with useful input from S.W. and K.M.Y. S.W. constructed the negative control *Btbd7* CRISPR/Cas9 knockouts. J.W.C. performed the bioinformatics and data analysis with useful input from D.M. D.M. performed the RNA-seq quality control analysis. G.C.B.C. performed the RNA-seq. J.W.C. wrote the manuscript with useful input from S.W. and K.M.Y.

## FUNDING

This work was supported by the National Institutes of Health Intramural Research Program, National Institute of Dental and Craniofacial Research [Z01 DE000524, Z01 DE000525 to K.M.Y.]; in part by the NIDCR Genomics and Computational Biology Core [ZIC DC000086]; and in part by the NIDCR Imaging Core [ZIC DE000750-01]. Funding for open access charge: National Institutes of Health.

## CONFLICT OF INTEREST

The authors declare they have no conflicts of interest.

## Notes

https://www.ncbi.nlm.nih.gov/geo/query/acc.cgi?acc=gse165689

https://github.com/collinsjw/Zcchc8-KO-pervasive-transcripts

### Competing Interest Statement

The authors have declared no competing interest.

### Summary of Updates

Updated Acknowledgments to include all members of Genomics and Computational Biology Core.

