## Supplementary figures and images for "ZCCHC8 is required for the degradation of pervasive transcripts originating from multiple genomic regulatory features"

### Graphical Abstract

#### GRAPHICAL ABSTRACT

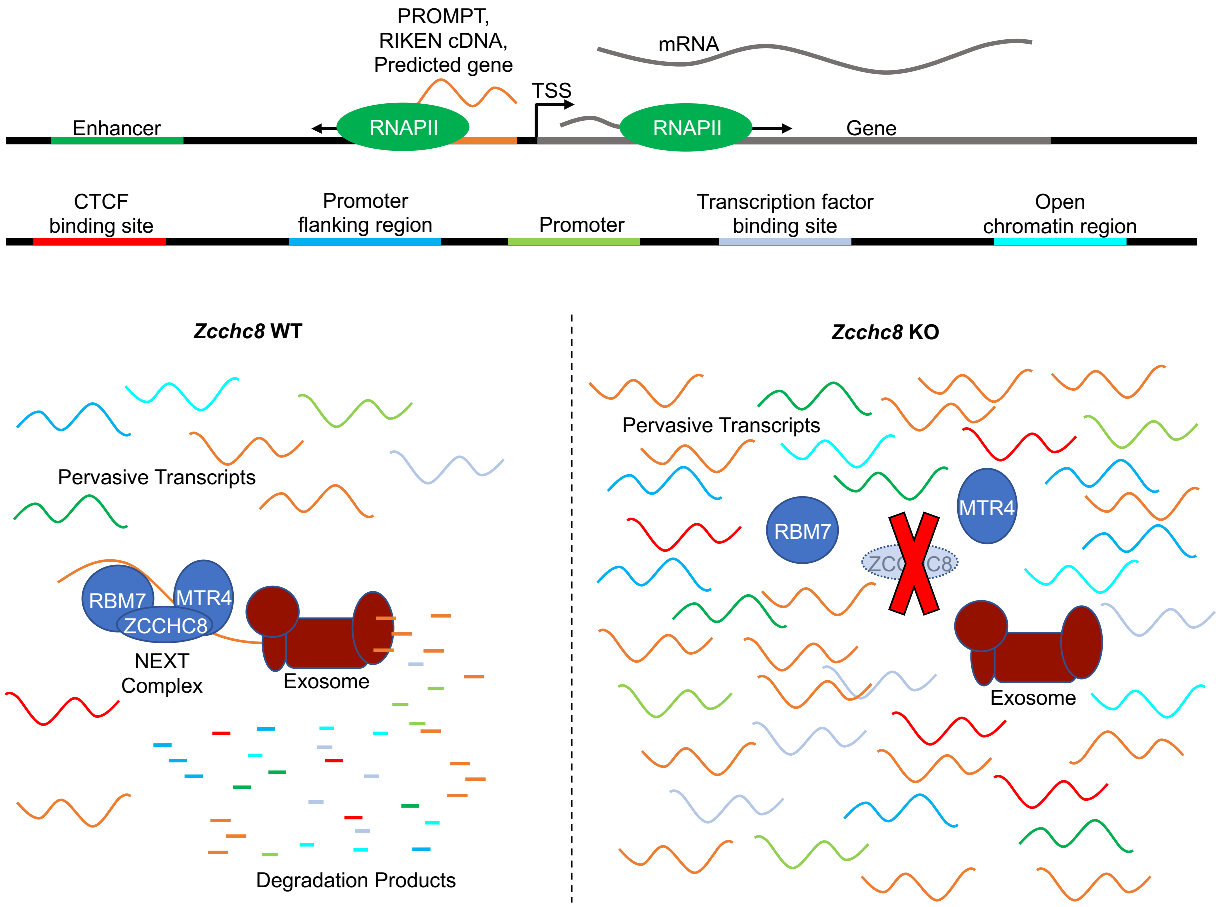
