## Supplemental Figure 1 for "ZCCHC8 is required for the degradation of pervasive transcripts originating from multiple genomic regulatory features"

### Figure S1

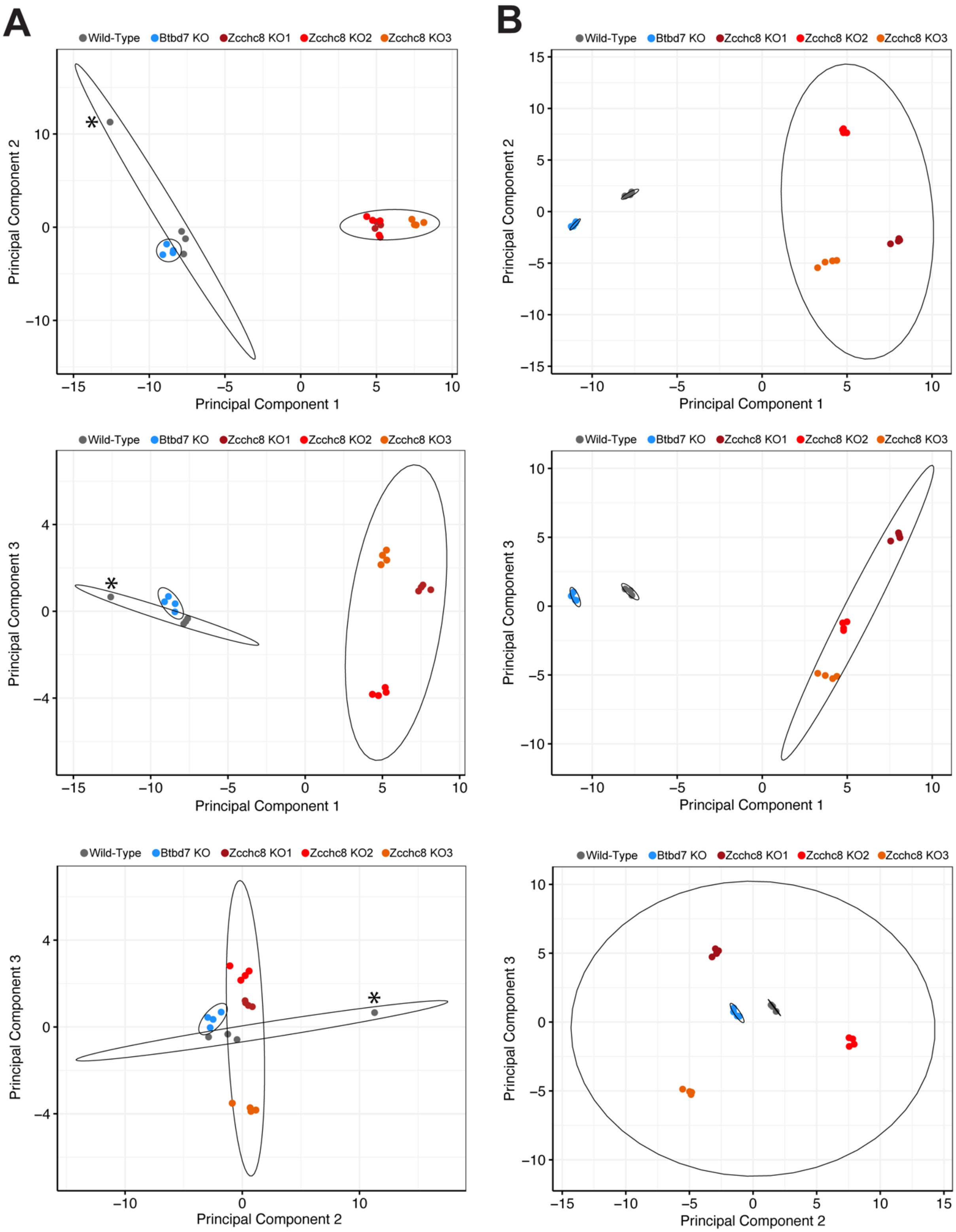
A
B
Wild-Type
Btbd7 KO
Zcchc8 KO1
Zcchc8 KO2
Zcchc8 KO3
Principal Component 1
Principal Component 2
Principal Component 3
*

Figure S1. (A) Principal component analysis plots depicting a single outlier sample from the wild-type group. Asterisks mark the outlier sample. Principal component variance: PC1 = 69.19%, PC2 = 10.81%, PC3 = 5.26%. (B) Principal component analysis plots after removal of the outlier sample from analysis. Principal component variance: PC1 = 58.41%, PC2 = 20.08%, PC3 = 11.61%. Ellipses represent 95% confidence regions in both (A) and (B).
