## Supplemental Figure 2 for "ZCCHC8 is required for the degradation of pervasive transcripts originating from multiple genomic regulatory features"

### Figure S2

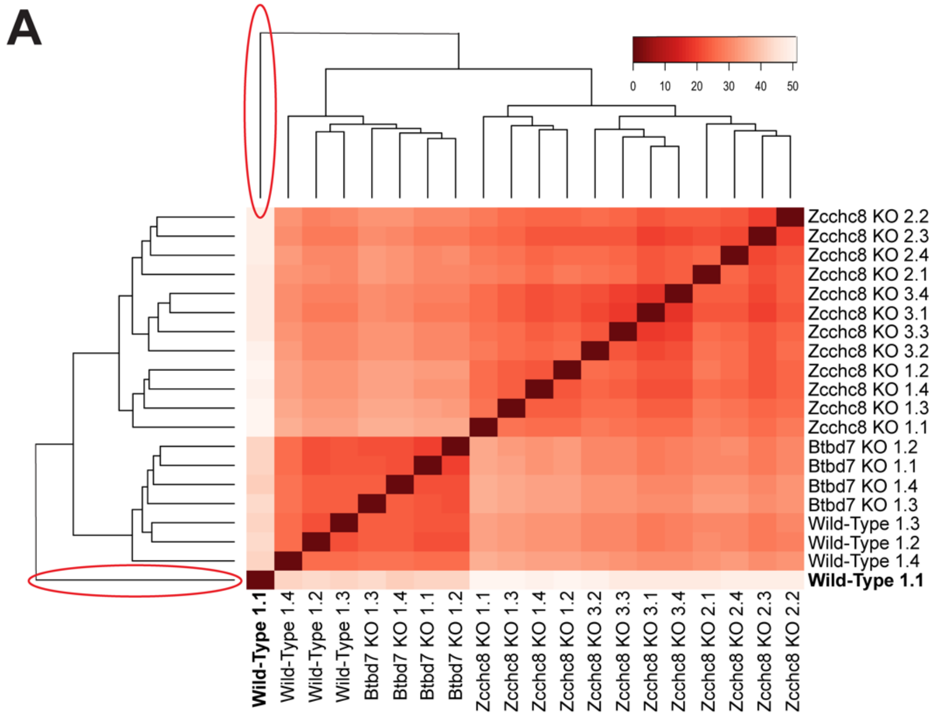
A
0
10
20
30
40
50
Zcchc8 KO 2.2
Zcchc8 KO 2.3
Zcchc8 KO 2.4
Zcchc8 KO 2.1
Zcchc8 KO 3.4
Zcchc8 KO 3.1
Zcchc8 KO 3.3
Zcchc8 KO 3.2
Zcchc8 KO 1.2
Zcchc8 KO 1.4
Zcchc8 KO 1.3
Zcchc8 KO 1.1
Btbd7 KO 1.2
Btbd7 KO 1.1
Btbd7 KO 1.4
Btbd7 KO 1.3
Wild-Type 1.3
Wild-Type 1.2
Wild-Type 1.4
Wild-Type 1.1
Wild-Type 1.1
Wild-Type 1.4
Wild-Type 1.2
Wild-Type 1.3
Btbd7 KO 1.3
Btbd7 KO 1.4
Btbd7 KO 1.1
Btbd7 KO 1.2
Zcchc8 KO 1.1
Zcchc8 KO 1.3
Zcchc8 KO 1.4
Zcchc8 KO 1.2
Zcchc8 KO 3.2
Zcchc8 KO 3.3
Zcchc8 KO 3.1
Zcchc8 KO 3.4
Zcchc8 KO 2.1
Zcchc8 KO 2.4
Zcchc8 KO 2.3
Zcchc8 KO 2.2

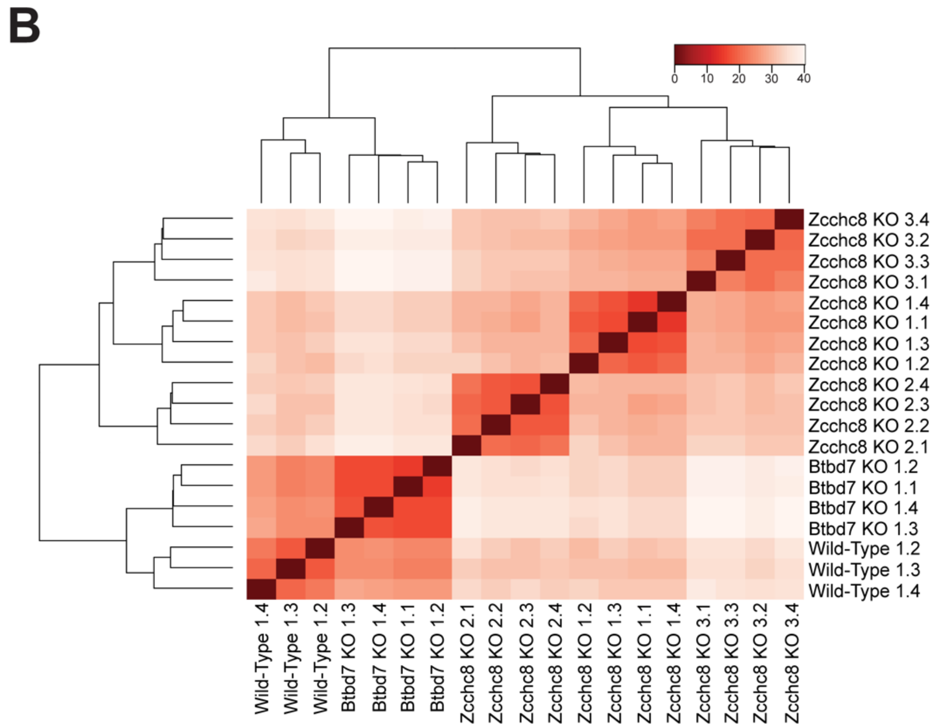
B
0
10
20
30
40
Zcchc8 KO 3.4
Zcchc8 KO 3.2
Zcchc8 KO 3.3
Zcchc8 KO 3.1
Zcchc8 KO 1.4
Zcchc8 KO 1.1
Zcchc8 KO 1.3
Zcchc8 KO 1.2
Zcchc8 KO 2.4
Zcchc8 KO 2.3
Zcchc8 KO 2.2
Zcchc8 KO 2.1
Btbd7 KO 1.2
Btbd7 KO 1.1
Btbd7 KO 1.4
Btbd7 KO 1.3
Wild-Type 1.2
Wild-Type 1.3
Wild-Type 1.4
Wild-Type 1.4
Wild-Type 1.3
Wild-Type 1.2
Btbd7 KO 1.3
Btbd7 KO 1.4
Btbd7 KO 1.1
Btbd7 KO 1.2
Zcchc8 KO 2.1
Zcchc8 KO 2.2
Zcchc8 KO 2.3
Zcchc8 KO 2.4
Zcchc8 KO 1.2
Zcchc8 KO 1.3
Zcchc8 KO 1.1
Zcchc8 KO 1.4
Zcchc8 KO 3.1
Zcchc8 KO 3.3
Zcchc8 KO 3.2
Zcchc8 KO 3.4

Figure S2. (A) Heatmap of sample-to-sample distances after unsupervised clustering of variance-stabilizing-transformed data. The wild-type 1.1 sample clustered individually on a single node (red circles, bold-type). (B) Heatmap produced using the same data analysis as in (A) after removal of the outlier wild-type 1.1 sample.
