## Supplemental Figure 3 for "ZCCHC8 is required for the degradation of pervasive transcripts originating from multiple genomic regulatory features"

**Figure S3**

A

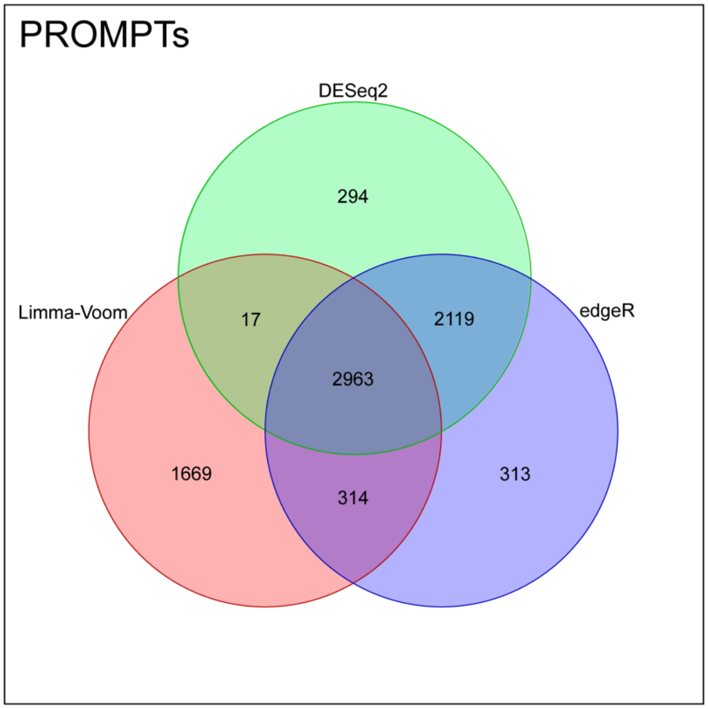
PROMPTs
DESeq2
294
Limma-Voom
17
2119
edgeR
2963
1669
314
313

B

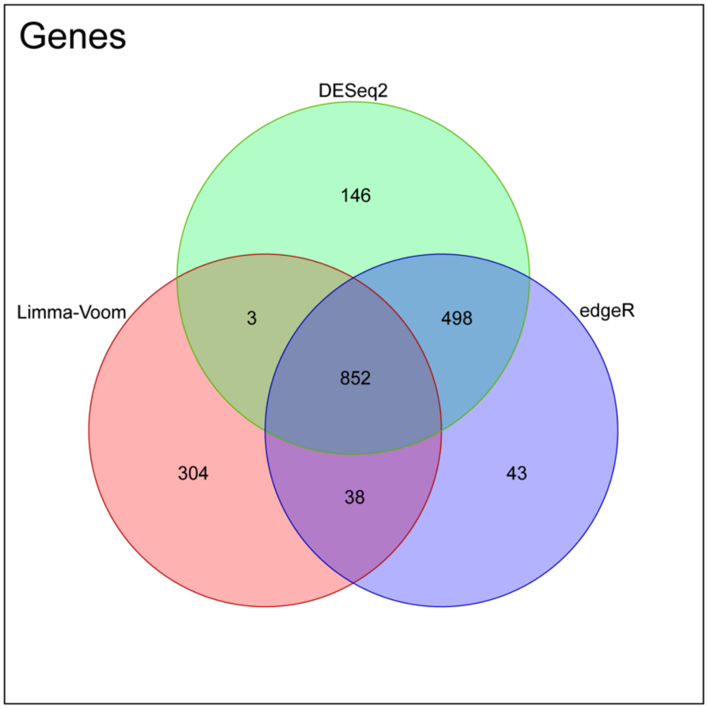
Genes
DESeq2
146
Limma-Voom
3
498
edgeR
852
304
38
43

Figure S3. (A) Venn diagram showing the number of differentially expressed PROMPTs determined via DESeq2, edgeR, and LimmaVoom analysis in SIMS *Zcchc8* knockout cells. (B) Venn diagram showing the number of differentially expressed genes determined via DESeq2, edgeR, and LimmaVoom analysis in SIMS *Zcchc8* knockout cells.
