## Supplemental Figure 4 for "ZCCHC8 is required for the degradation of pervasive transcripts originating from multiple genomic regulatory features"

### Figure S4

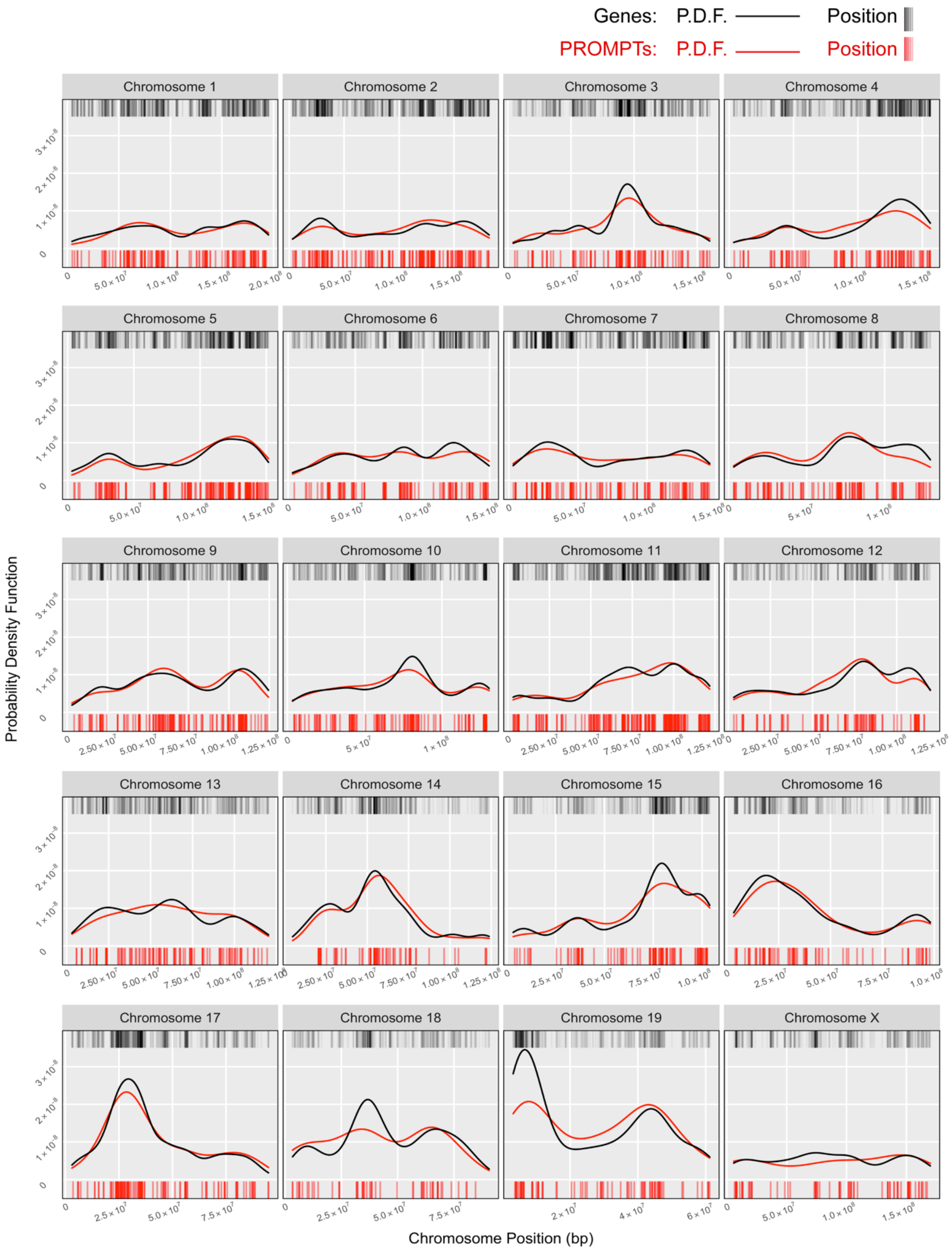
Genes: P.D.F. Position
PROMPTs: P.D.F. Position
Chromosome 1
Chromosome 2
Chromosome 3
Chromosome 4
Chromosome 5
Chromosome 6
Chromosome 7
Chromosome 8
Chromosome 9
Chromosome 10
Chromosome 11
Chromosome 12
Chromosome 13
Chromosome 14
Chromosome 15
Chromosome 16
Chromosome 17
Chromosome 18
Chromosome 19
Chromosome X
Probability Density Function
Chromosome Position (bp)

Figure S4. Probability density function and rug plots indicating the distribution of PROMPTs closely matches the distribution of expressed genes along each chromosome. Rug plots mark individual genes (black) and PROMPTs (red) as vertical tick marks above and below, respectively. Probability density function curves for genes (black) and PROMPTs (red) were generated using the Gaussian kernel density estimate with Silverman's rule of thumb bandwidth method (Nrd0).
