## Supplemental Figure 5 for "ZCCHC8 is required for the degradation of pervasive transcripts originating from multiple genomic regulatory features"

### Figure S5

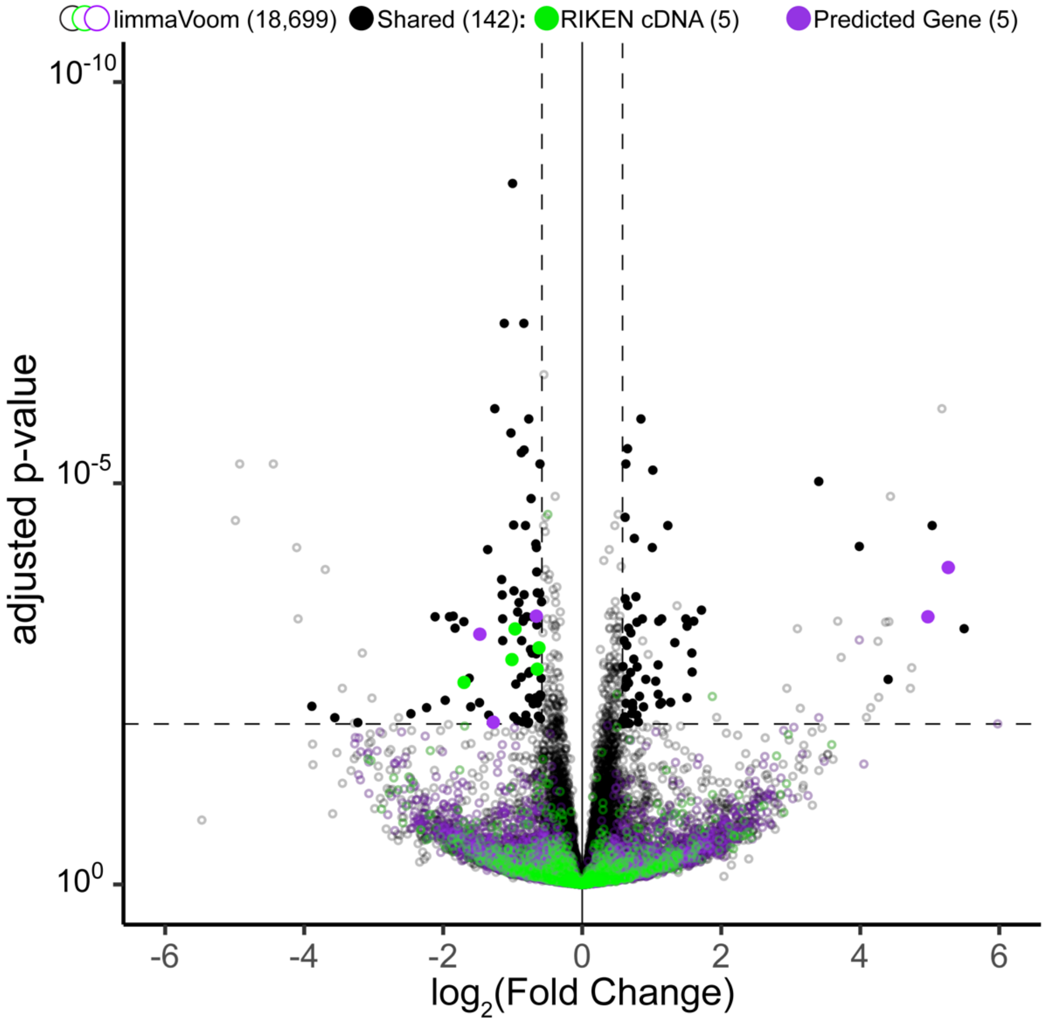
limmaVoom (18,699)
Shared (142):
RIKEN cDNA (5)
Predicted Gene (5)
adjusted p-value
$10^{-10}$
$10^{-5}$
$10^{0}$
-6
-4
-2
0
2
4
6
$log_2$(Fold Change)

Figure S5. Scatter plot of gene expression data in SIMS *Btbd7* knockout cells. Data were generated using the LimmaVoom statistical analysis. Horizontal and vertical dashed lines demarcate adjusted p-value of 0.01 and fold-change of 1.5 ($\log_2(1.5) \approx 0.584$), respectively. Open circles indicate those genes that are specific to the LimmaVoom analysis. Closed circles indicate genes that are shared within DESeq2, edgeR, and LimmaVoom analyses and meet the significance thresholds of >1.5 fold-change and <0.01 adjusted p-value. Colored circles indicate RIKEN cDNAs (green) and predicted genes (magenta).
