## Supplemental Table 1 for "ZCCHC8 is required for the degradation of pervasive transcripts originating from multiple genomic regulatory features"

**Table S1.** Differentially expressed genomic regulatory features and genes by statistical method

| | DESeq2 | | edgeR | | LimmaVoom | | Shared | |
|---|---|---|---|---|---|---|---|---|
| | Up | Down | Up | Down | Up | Down | Up | Down |
| **PROMPTs** | | | | | | | | |
| *Zcchc8* KO | 4618 | 775 | 4353 | 1356 | 2533 | 2430 | 2486 | 477 |
| *Btbd7* KO | 42 | 26 | 40 | 36 | 10 | 12 | 5 | 8 |
| **Enhancers** | | | | | | | | |
| *Zcchc8* KO | 439 | 50 | 400 | 78 | 258 | 68 | 243 | 28 |
| *Btbd7* KO | 9 | 4 | 10 | 6 | 11 | 5 | 3 | 3 |
| **Promoters** | | | | | | | | |
| *Zcchc8* KO | 338 | 42 | 327 | 63 | 112 | 38 | 105 | 22 |
| *Btbd7* KO | 6 | 3 | 8 | 7 | 4 | 2 | 1 | 2 |
| **PFR** | | | | | | | | |
| *Zcchc8* KO | 839 | 75 | 771 | 126 | 487 | 157 | 471 | 51 |
| *Btbd7* KO | 17 | 11 | 14 | 6 | 13 | 8 | 2 | 3 |
| **CTCF sites** | | | | | | | | |
| *Zcchc8* KO | 469 | 38 | 425 | 67 | 269 | 71 | 247 | 19 |
| *Btbd7* KO | 9 | 4 | 4 | 2 | 9 | 9 | 2 | 2 |
| **TFBS** | | | | | | | | |
| *Zcchc8* KO | 86 | 10 | 72 | 13 | 48 | 15 | 45 | 6 |
| *Btbd7* KO | 3 | 2 | 3 | 0 | 1 | 0 | 1 | 0 |
| **OCR** | | | | | | | | |
| *Zcchc8* KO | 239 | 27 | 224 | 35 | 151 | 48 | 142 | 9 |
| *Btbd7* KO | 3 | 4 | 1 | 4 | 3 | 1 | 0 | 0 |
| **Genes** | | | | | | | | |
| *Zcchc8* KO | 1342 | 157 | 1211 | 220 | 728 | 469 | 712 | 140 |
| *Btbd7* KO | 95 | 131 | 100 | 138 | 85 | 91 | 62 | 80 |

Table S1. Differentially expressed genomic regulatory features and genes as determined by three different analyses using cut-off values of >1.5-fold expression difference and adjusted p-value <0.01. The number of features that were commonly found within all three statistical analyses are indicated in the Shared column. PFR = Promoter Flanking Regions, TFBS = Transcription Factor Binding Sites, OCR = Open Chromatin Regions.
