## Supplemental Table 2 for "ZCCHC8 is required for the degradation of pervasive transcripts originating from multiple genomic regulatory features"

**Table S2.** Gene expression of RNA degradation complex subunits in *Zcchc8* KO cells

| | DESeq2 | | edgeR | | LimmaVoom | |
|---|---|---|---|---|---|---|
| | $log_2$(FC) | Adj. p-value | $log_2$(FC) | Adj. p-value | $log_2$(FC) | Adj. p-value |
| **NEXT** | | | | | | |
| *Mtr4* | −0.26 | 0.013 | −0.28 | 0.015 | −0.20 | 0.075 |
| *Rbm7* | −0.12 | 0.371 | −0.13 | 0.354 | −0.08 | 0.503 |
| **PAXT** | | | | | | |
| *Pabpn1* | 0.25 | 0.347 | 0.23 | 0.442 | 0.16 | 0.532 |
| *Rbm26* | −0.17 | 0.029 | −0.18 | 0.063 | −0.12 | 0.146 |
| *Rbm27* | 0.12 | 0.231 | 0.10 | 0.419 | 0.17 | 0.031 |
| *Zc3h3* | 0.18 | 0.703 | 0.16 | 0.789 | −0.04 | 0.923 |
| *Zfc3h1* | 0.14 | 0.231 | 0.12 | 0.352 | 0.20 | 0.045 |
| **RNA Exosome** | | | | | | |
| *Dis3* | −0.31 | 0.0006 | −0.33 | 0.001 | −0.26 | 0.011 |
| *Dis3l* | −0.24 | 0.195 | −0.26 | 0.163 | −0.26 | 0.119 |
| *Exosc1* | −0.37 | 0.0004 | −0.39 | 0.001 | −0.34 | 0.004 |
| *Exosc2* | 0.04 | 0.903 | 0.02 | 0.968 | −0.02 | 0.937 |
| *Exosc3* | 0.21 | 0.430 | 0.19 | 0.512 | 0.04 | 0.894 |
| *Exosc4* | −0.27 | 0.048 | −0.28 | 0.062 | −0.30 | 0.019 |
| *Exosc5* | 0.20 | 0.312 | 0.19 | 0.432 | 0.14 | 0.468 |
| *Exosc7* | −0.18 | 0.567 | −0.20 | 0.513 | −0.19 | 0.424 |
| *Exosc8* | −0.35 | 0.0009 | −0.37 | 0.002 | −0.32 | 0.006 |
| *Exosc9* | −0.40 | 0.014 | −0.41 | 0.015 | −0.41 | 0.011 |
| *Exosc10* | −0.06 | 0.651 | −0.07 | 0.598 | −0.02 | 0.886 |

Table S2. Gene expression data for RNA degradation complex subunits as computed using three different statistical analyses: DESeq2, edgeR, and LimmaVoom. Adj. p-value = adjusted p-value; FC = fold-change.
