## Supplemental Table 3 for "ZCCHC8 is required for the degradation of pervasive transcripts originating from multiple genomic regulatory features"

**Table S3.** Intersections of differentially expressed genomic regulatory features and genes by dataset

| | SIMS | | E12.5 Brains | | ES Cells | | SIMS/ E12.5 Brains | | SIMS/ ES Cells | | SIMS/ E12.5 Brains/ ES Cells | |
|---|---|---|---|---|---|---|---|---|---|---|---|---|
| | Up | Down | Up | Down | Up | Down | Up | Down | Up | Down | Up | Down |
| **PROMPTs** | 2486 | 477 | 4740 | 746 | 422 | 300 | 1772 | 26 | 138 | 24 | 132 | 1 |
| **Enhancers** | 243 | 28 | 775 | 12 | 152 | 47 | 119 | 0 | 7 | 0 | 7 | 0 |
| **Promoters** | 105 | 22 | 438 | 25 | 143 | 68 | 50 | 0 | 6 | 1 | 5 | 0 |
| **PFR** | 471 | 51 | 1342 | 40 | 235 | 86 | 221 | 0 | 19 | 1 | 16 | 0 |
| **CTCF sites** | 247 | 19 | 1159 | 14 | 101 | 27 | 139 | 0 | 5 | 1 | 5 | 0 |
| **TFBS** | 45 | 6 | 198 | 1 | 21 | 9 | 22 | 0 | 0 | 0 | 0 | 0 |
| **OCR** | 142 | 9 | 638 | 7 | 61 | 17 | 71 | 0 | 2 | 0 | 2 | 0 |
| **Genes** | 712 | 140 | 1410 | 131 | 1804 | 1297 | 450 | 1 | 89 | 7 | 69 | 0 |

Table S3. Differentially expressed genomic regulatory features and genes as determined by three different analyses using cut-off values of >1.5-fold expression difference and adjusted p-value <0.01. The number of features that were commonly found in SIMS *Zcchc8* knockout cells and E12.5 brains (GSE126108) or ES cells (GSE127790) or all three are indicated in the last three columns. PFR = Promoter Flanking Regions, TFBS = Transcription Factor Binding Sites, OCR = Open Chromatin Regions.
